# Ventral subiculum inputs to nucleus accumbens medial shell preferentially innervate D2R medium spiny neurons and contain calcium permeable AMPARs

**DOI:** 10.1101/2022.10.12.511974

**Authors:** Emma E Boxer, JungMin Kim, Brett Dunn, Jason Aoto

**Affiliations:** University of Colorado Anschutz Medical Campus, Aurora, CO, 80045, USA

## Abstract

Ventral subiculum (vSUB) is the major output region of ventral hippocampus (vHIPP) and sends major projections to nucleus accumbens medial shell (NAcMS). Hyperactivity of the vSUB-NAcMS circuit is associated with substance use disorders (SUDs) and the modulation of vSUB activity alters drug seeking and drug reinstatement behavior in rodents. However, to the best of our knowledge, the cell-type specific connectivity and synaptic transmission properties of the vSUB-NAcMS circuit have never been directly examined. Instead, previous functional studies have focused on total ventral hippocampal (vHIPP) output to NAcMS without distinguishing vSUB from other subregions of vHIPP, including ventral CA1 (vCA1). Using *ex vivo* electrophysiology, we systematically characterized the vSUB-NAcMS circuit with cell-type and synapse specific resolution in male and female mice and found that vSUB output to dopamine receptor type-1 (D1R) and type-2 (D2R) expressing medium spiny neurons (MSNs) displays a functional connectivity bias for D2R MSNs. Furthermore, we found that vSUB-D1R and -D2R MSN synapses contain calcium-permeable AMPA receptors in drug-naïve mice. Finally, we find that, distinct from other glutamatergic inputs, cocaine exposure selectively induces plasticity at vSUB-D2R synapses. Importantly, we directly compared vSUB and vCA1 output to NAcMS and found that vSUB synapses are functionally distinct and that vCA1 output recapitulated the synaptic properties previously ascribed to vHIPP. Our work highlights the need to consider the contributions of individual subregions of vHIPP to SUDs and represents an important first step toward understanding how the vSUB-NAcMS circuit contributes to the etiologies that underlie SUDs.

## Introduction

The nucleus accumbens medial shell (NAcMS) plays critical roles in reward and motivated behavior. NAc is primarily populated by medium spiny principal neurons (MSNs) that are defined by their expression of either dopamine receptor type 1 (D1R) or type 2 (D2R). The respective roles of D1R and D2R MSNs in drug seeking behavior has been contentious – the predominant model contends that D1R MSNs promote reward seeking whereas D2R MSNs inhibit it (Lobo et al., 2010; Lobo & Nestler, 2011; Bock et al., 2013). However, several studies point to a more complicated role for MSNs in drug seeking that appears to be dependent on the subregion of NAc studied (Cole et al., 2018; Z. Liu et al., 2022; Soares-Cunha et al., 2022, 2016). In dorsal NAcMS, activation of D2R MSNs promotes reward, a behavior presumably mediated via their unique connections to glutamatergic neurons in ventral pallidum (VP) (Yao et al., 2021). Thus, it is critical to dissect MSN function within the context of discrete NAc subregions that are defined by their anatomical position and by their input and output circuitry, however, an understanding of the inputs that drive D2R MSN activity in this region is incomplete (Baimel et al., 2019; Castro & Bruchas, 2019; Yang et al., 2018; Yao et al., 2021).

The ventral hippocampus (vHIPP) provides the most robust source of glutamatergic input to NAcMS (Britt et al., 2012; Li et al., 2018). This excitatory circuit regulates drug-relevant locomotion and cue and context-induced reinstatement of drug seeking in rodents (Bossert et al., 2016; Britt et al., 2012; Glangetas et al., 2015; Grace, 2010; Marchant et al., 2016; Preston et al., 2019). vHIPP input displays biased synaptic strength onto D1R MSNs (MacAskill et al., 2014), and long-term potentiation induced by withdrawal from cocaine exposure has been observed at vHIPP-D1R MSN synapses, but not D2R MSNs (Pascoli et al., 2014). Importantly, vHIPP is a composite of afferents originating from functionally discrete subregions of vHIPP: ventral CA1 (vCA1) and ventral subiculum (vSUB). vSUB is the major output of vHIPP and sends robust projections to dorsal NAcMS (Lopes da Silva et al., 1984; Britt et al., 2012; Li et al., 2018; Yao et al., 2021). Further, vSUB is thought to be a key regulator of reward seeking behavior as the selective pharmacological manipulation of vSUB activity is sufficient to alter drug reinstatement in rodents (Bossert et al., 2016; Marchant et al., 2016). Distinct from vCA1, vSUB comprises two classes of excitatory principal neurons identified electrophysiologically as regular or burst spiking (RS or BS) cells. RS and BS neurons are equally represented in vSUB but exhibit distinct gene expression, connectivity, plasticity, and behavioral functions (Wozny et al., 2008; Cembrowski et al., 2018; Boxer et al., 2021; Böhm et al., 2015; Graves et al., 2012). Thus, RS and BS vSUB neurons may provide distinct input to dorsal NAcMS and possess unique synaptic properties compared to other subregions of vHIPP. While most studies have examined the synaptic properties of total vHIPP or vCA1 input to NAc, to our knowledge, the functional connectivity and synaptic properties of vSUB input to dorsal NAcMS are untested.

Here, we systematically dissected the connectivity and functional properties of the vSUB-NAcMS circuit and reveal unique qualities of vSUB input. Despite a disproportionate amount of RS neurons projecting to total nucleus accumbens shell (NAcSh), we performed the first intersectional cell-type specific retrograde labeling of the vSUB-NAcMS circuit and found that RS and BS neurons project equally to D1R and D2R MSNs. We next functionally characterized vSUB input in dorsal NAcMS. First, vSUB displays a significant bias for D2R MSNs. Second, vSUB synapses onto both MSNs in drug naïve animals unexpectedly contain calcium-permeable AMPA receptors (CP-AMPARs). Third, cocaine administration selectively potentiates vSUB-D2R but not vSUB-D1R MSNs. Our findings provide critical insight into the unique synaptic properties of vSUB input to dorsal NAcMS D2R MSNs.

## Results

### Cell-type specific circuit tracing

The cell-type-specific connectivity of RS and BS neurons in vSUB with D1R vs D2R MSNs in NAcMS, has not been fully explored (Figure 1A-B). We first aimed to confirm previous findings by assessing the identity of presynaptic vSUB neurons that generally project to the NAcMS. Using wild-type mice, we stereotactically injected mRuby expressing retrograde AAV_2_ (AAV_2_rg-mRuby) into NAcMS (Figure 1C), then made *ex vivo* slices and performed whole cell patch clamp to electrophysiologically characterize mRuby^+^ neurons in vSUB. Consistent with previous reports, mRuby^+^ NAcMS-projecting neurons were concentrated in proximal vSUB but also present in ventral entorhinal cortex (vEC), ventral CA1 (vCA1) (Naber & Witter, 1998; Britt et al., 2012; Cembrowski et al., 2018; Lee et al., 2019; Williams et al., 2020). Although vSUB is composed of about 50% RS and 50% BS neurons (Staff et al., 2000; Graves et al., 2012), we found that the majority of NAcMS-projecting neurons in vSUB were RS neurons (70% ± 4) (Figure 1G), which is remarkably consistent with other studies using different retrograde labeling approaches such as retrobeads or cholera toxin B (Kim & Spruston, 2012; Lee et al., 2019). In light of our recent findings that the wiring of vSUB local circuitry is sexually dimorphic (Boxer et al., 2021), we reviewed whether the wiring of the vSUB-NAcMS circuit also exhibits sexual dimorphism. We separated our AAV_2_rg-mRuby experiment by sex but found that equivalent proportions of RS/BS neurons project to NAcMS in male and female mice (%RS: female: 66 ± 4, male: 75 ± 6; U = 2, p = 0.40, Mann-Whitney) (Figure 1G, females: closed circles, males: open circles).

**Figure 1.**
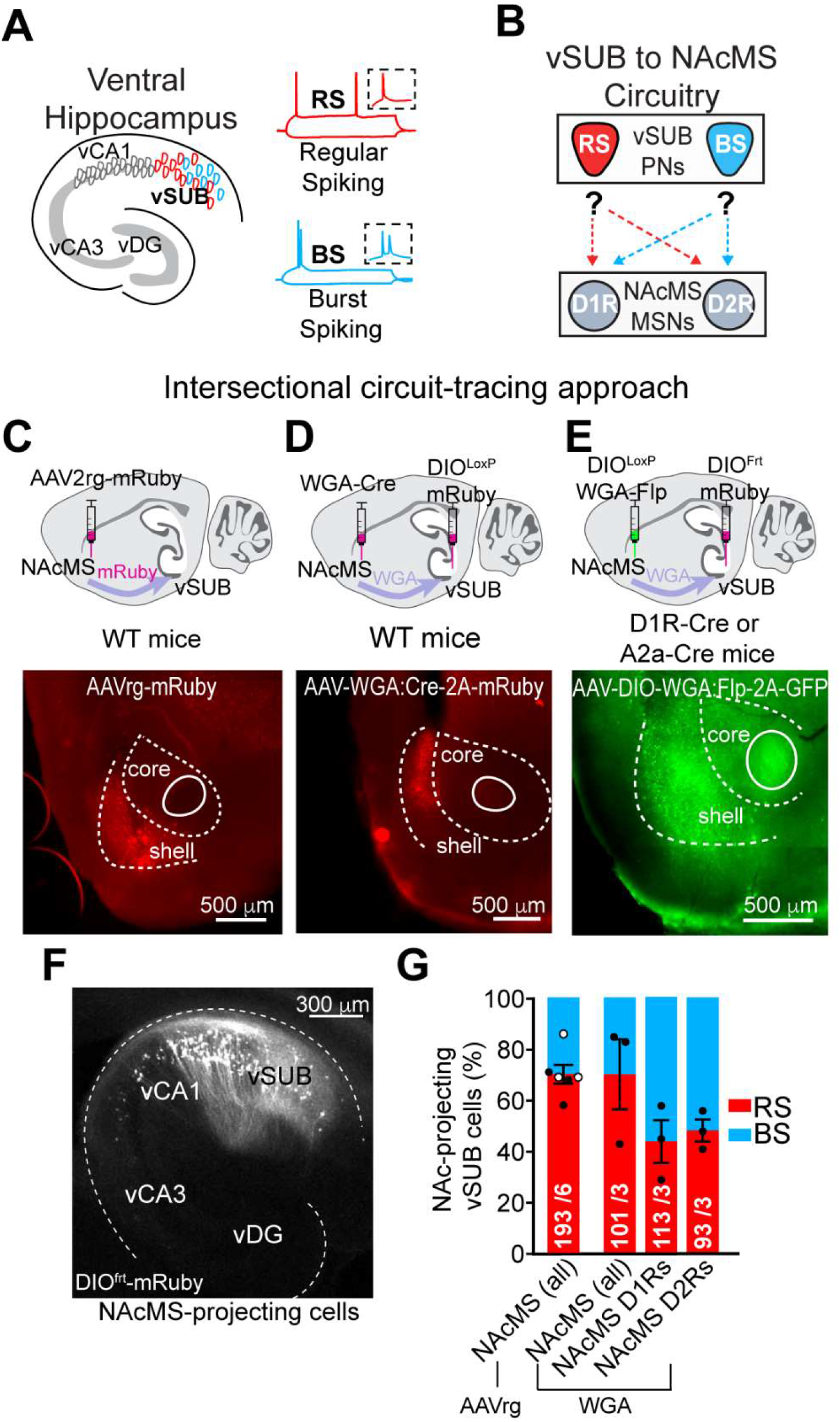
vSUB regular and burst spiking neurons equally project to NAc medial shell D1R and D2R MSNs. (A) Left: Schematic illustration of vHIPP in the horizontal plane. Ventral subiculum (vSUB) mainly consists of 2 distinct principal neuron types -regular and burst spiking neurons (RS and BS). Right: Representative current-clamp traces of RS (above) and BS (below) principal neurons. Insets: Magnification of the first action potential in response to current injection. (B) Simplified graphic of vSUB to NAcMS circuitry. The proportion of RS and BS principal neurons in vSUB projecting to D1R and D2R MSNs in NAcSh is unknown. (C-E) Above: Schematic drawing of intersectional circuit-tracing approaches and mouse lines used for characterizing projection pattern of vSUB-NAcMS circuit. Below: Representative image of viral expression in NAcMS in coronal slices. Scale bar 500 μm. (C) AAV-Retro-mRuby was injected in NAcMS of WT mice, labeling all NAcMS projecting cells. (D) WGA:Cre-recombinase is retrogradely transported from NAcMS to vSUB output neurons that are labeled by injection of Cre-dependent mRuby to label all NAcMS-projecting vSUB cells. (E) Cre-dependent WGA:Flp-recombinase is selectively expressed D1R or D2R MSN populations in NAcMS and retrogradely transported to vSUB output neurons that express Flp-dependent mRuby to permit visualization and identification of D1R or D2R MSN projecting vSUB neurons. (F) Representative image of a horizontal vHIPP slice after intersectional labeling using AAV-DIO-WGA:Flp-2A-GFP with AAV-DIO^frt^-mRuby shows labeling of NAcMS D1R MSN-projecting cells is largely restricted to vSUB. Scale bar: 300 μm. (G) Summary graph of RS and BS principal neurons projecting to NAcMS using different viral approaches and transgenic mouse lines. Independent of postsynaptic cell-type, RS neurons send more projections to overall NAcMS than BS neurons (left two bars). There is no difference in proportion of RS and BS neurons projecting to NAcMS between females and males (far left bar; females: closed circles, males: open circles). Similar proportions of RS and BS neurons project to D1R and D2R MSNs. Number of cells recorded and animals used for each method is included in the graph. Error bars ± SEM. RS: regular spiking, BS: burst spiking, MSN: medium spiny neuron, vHIPP: ventral hippocampus, vSUB: ventral subiculum, NAcMS: nucleus accumbens shell, NAcMS: nucleus accumbens medial shell.

Given the strikingly disproportionate level of vSUB RS neuron input to NAcMS, we asked if RS neurons also disproportionally innervate D1R and D2R MSNs. To address this question, we employed an intersectional viral circuit tracing approach utilizing wheat germ agglutinin (WGA), which functions as a transcellular retrograde tracer, and D1R-Cre and A2a-Cre mouse lines to restrict expression of WGA to either D1R or D2R MSNs, respectively (Figure 1E) (Gradinaru et al., 2010). We stereotactically co-injected AAVs carrying Cre-dependent WGA fused to Flp recombinase (DIO^LoxP^-WGA-Flp) into NAcMS, and injected AAVs carrying Flp-dependent mRuby (DIO^Frt^-mRuby) into vSUB (Figure 1E). WGA-Flp, selectively expressed in either D1R or D2R MSNs, is retrogradely transported into vSUB, where it selectively enables the expression of Flp-dependent mRuby in vSUB neurons that innervate D1R or D2R MSNs (Figure 1F). mRuby^+^ neurons were then identified electrophysiologically in *ex vivo* slices. Using this intersectional approach, we found that vSUB RS and BS neurons surprisingly projected in roughly equal proportions to both D1R and D2R MSNs; RS neurons represented ∼45% of total vSUB input to each cell type (Figure 1G).

Intriguingly, this is in contrast to retrograde labeling of *total* NAcMS that identified RS neurons as the dominant source of input. This suggests that the RS neurons that represent the major input to *total* NAcMS, which includes MSNs as well as local GABAergic and cholinergic interneurons, may primarily target non-MSN cell populations. To test if this unexpected cell-type specific projection pattern was due to inherent differences between AAVrg- and WGA-mediated circuit tracing approaches, we injected cre-independent WGA-Flp into NAcMS and Flp-dependent mRuby into vSUB of wild-type animals (Figure 1D). As expected, we found that RS neurons represented ∼70% of the NAcMS-projecting vSUB population, consistent with our AAVrg results and with previous retrograde tracing experiments (Figure 1G). Thus, the proportions of RS and BS neurons that provide input onto D1R and D2R MSNs, systematically identified by intersectional circuit tracing, are not artifacts of WGA uptake efficiency, but are likely a faithful representation of vSUB-MSN cell-type specific connectivity. Thus, our results indicate that a significant amount of vSUB output is not to MSNs, but to other cell types in NAcMS, which includes parvalbumin (PV) and somatostatin GABAergic interneurons and cholinergic interneurons. This is consistent with the finding that vHIPP excitatory synaptic strength is significantly greater to PV-interneurons in NAcSh than to MSNs (Scudder et al., 2018; Yu et al., 2017). Our findings also show that BS neurons are more highly represented in the vSUB to NAcMS MSN circuit than the vSUB to NAcMS interneuron circuit.

### vSUB input to NAcMS is biased to D2R MSNs

The properties of vSUB synapses on D1R and D2R MSNs in NAcMS have not been directly assessed and may be distinct from other areas of the extended ventral hippocampal formation which also project to NAc, such as vCA1 and vEC (Okuyama et al., 2016). To examine the specific functional connectivity of vSUB-D1R and vSUB-D2R MSN synapses in NAcMS, we stereotactically injected vSUB of D1R-tdTomato mice with AAV-ChIEF-mRuby on P21 (Figure 2A). We allowed ChIEF-mRuby 5-6 weeks to express before making *ex vivo* coronal slices of NAc. Immediately before the start of each experiment, we also generated serial horizontal brain sections, which permit precise examination of discrete hippocampal subfields, to confirm that ipsilateral vSUB injections had no, or very minimal, viral spread to CA1 or EC (Figure 2B). Hemispheres with spillover to vCA1 or vEC were excluded from this study. We found that vSUB terminals were concentrated in the dorsal NAcMS (Figure 2B), consistent with what has been observed following total vHIPP injections of ChR2 (Britt et al., 2012; MacAskill et al., 2014). We then performed dual whole cell recordings in *ex vivo* slices of nearby TdTomato^+^ (D1R) and TdTomato^−^ (putative D2R) MSNs in dorsal NAcMS and performed terminal stimulation to measure light-evoked excitatory postsynaptic currents (EPSCs). As vSUB fiber density varies along the rostral-caudal axis of NAcMS (Groenewegen et al., 1999), recording from neighboring pairs of D1R and D2R MSNs allows for the direct comparison of vSUB input to D1R vs D2R MSNs. We first sought to characterize the relative strengths of vSUB-D1R and vSUB-D2R MSN excitatory synapses by assessing the input-output (I/O) relationship. As expected, given the robust input of vSUB to NAcMS, optogenetic stimulation of vSUB terminals produced reliable EPSCs in both classes of MSNs. Interestingly, we found that vSUB EPSC amplitudes at individual light intensities (D1R vs D2R at 1.8, 3.45, 5.78 mW, U = 78, 63, 47, p = 0.034, 0.008, 0.004, respectively, Mann-Whitney) (Figure 2C) and the corresponding slopes of the I/O curves were significantly stronger onto D2R MSNs compared to D1R MSNs (U = 40, p = 0.0003, Mann-Whitney) (Figure 2C).

**Figure 2.**
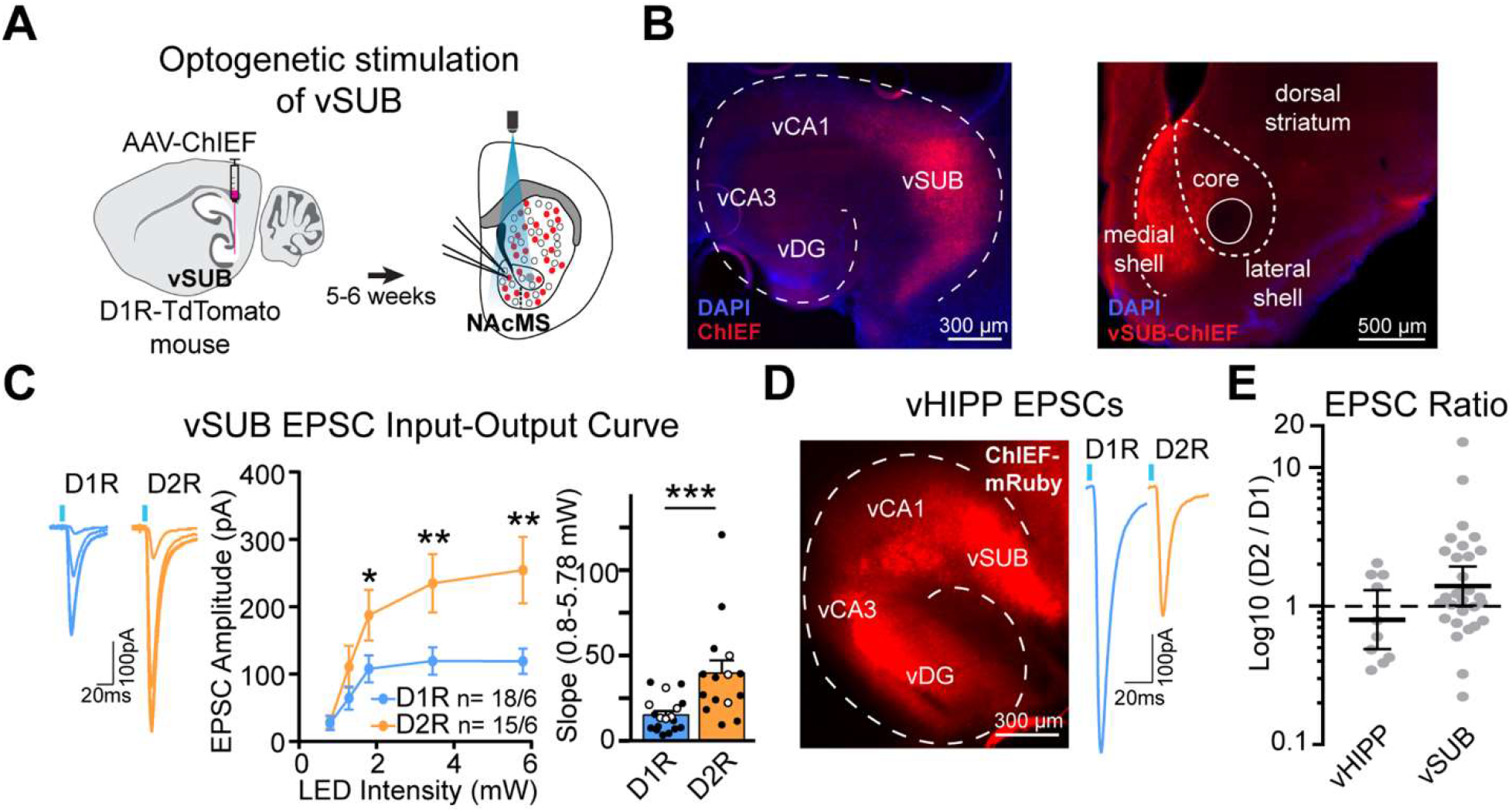
vSUB input to NAc medial shell is biased to D2R MSNs. (A) Experimental schematic to isolate vSUB output to D1R and D2R MSNs: AAV-ChIEF was injected in vSUB of ∼P21 D1R-TdTomato mouse. At P56-63, coronal NAc slices were made and vSUB input onto D1R and D2R MSNs in NAcMS was measured by optical stimulation of vSUB terminals. (B) Left: Representative image of ChIEF-mRuby injected vSUB in the horizontal plane. Scale bar: 300 μm. Right: Representative image of vSUB fibers infected with ChIEF-mRuby enriched in NAcMS of a coronal NAc slice. Scale bar: 500 μm. (C) D2R MSNs receive stronger input from vSUB than D1R MSNs in NAcMS. Representative light-evoked EPSCs (left), Input/Output (I/O) curves (middle) and corresponding summary graph of the I/O slope (right) of light-evoked EPSC amplitudes in D1R and D2R MSNs. There is no difference in slope of EPSC I/O in D1R and D2R MSNs between females and males. (I/O slope; females: closed circles, males: open circles) (D) Left: Representative image of ChIEF-mRuby expression in the entire vHIPP in horizontal plane. Scale bar: 300 μm. Right: Representative traces of light evoked EPSCs recorded from simultaneous dual recordings of D1R and D2R MSNs. (E) EPSC ratio of NAcMS D2R/D1R MSNs determined from optogenetic activation of vHIPP or vSUB terminals. Data are presented as geometric mean with 95% confidence intervals. Note logarithmic scale of Y-axis. vHIPP n=10 pairs from 2 mice; vSUB n=30 pairs from 14 mice. Error bars ± SEM. Number of cells and animals used for each experiment is included in each figure or corresponding figure legend.

The bias of vSUB-specific synaptic strength on D2R MSNs is unexpected because it diverges from previous findings from *total* vHIPP that revealed synaptic strength is biased for D1R MSNs (MacAskill et al., 2014). This could indicate that the functional organization of the vSUB projections in dorsal NAcMS is unique from the *total* vHIPP. Importantly, vHIPP output represents an amalgam of afferents originating from vCA1, vSUB, and possibly vEC. To directly compare the properties of synapses made by vHIPP and vSUB onto D1R and D2R MSNs, we infected entire vHIPP or vSUB with AAV-ChIEF (Figure 2D) and stimulated terminals in dorsal NAcMS while performing dual recordings from pairs of neighboring D1R and D2R MSNs. We calculated the D2R/D1R EPSC ratios to compare the relative cell-type specific synaptic strengths of each input. Consistent with the previously observed vHIPP-D1R bias, the D2/D1 ratio in vHIPP injected animals was <1, whereas in vSUB injected animals, the D2/D1 ratio was >1, which is consistent with a vSUB-D2R bias (geometric means: vHIPP = 0.797 (95% CI = 0.488 -1.3), vSUB = 1.394 (95% CI = 1.01 -1.92)) (Figure 2E). Our findings support the notion that the vSUB to NAcMS circuit possesses connectivity and synaptic properties unique from other hippocampal subfield inputs and argues that specific targeting of vSUB is necessary to reveal these subregion specific characteristics. This is further supported by a recent study that performed rabies-based circuit tracing experiments and found that more vSUB neurons project to D2R MSNs than to D1R MSNs in NAcSh, whereas vCA1 neurons exhibited no bias (Li et al., 2018)

Finally, the sexually dimorphic organization of vSUB local circuitry (Boxer et al., 2021) prompted us to ask whether the vSUB-specific bias for D2R MSNs was sex-specific. We separated our data by sex but found no sex differences (effect of sex: ns, effect of cell-type: F(1,29) = 6.913, p = 0.014, effect of interaction: ns, Ordinary Two-way ANOVA) (Figure 2C, females: closed circles, males: open circles). Thus, although RS and BS neurons in vSUB are under local sex-specific inhibitory control, their output to NAc MSNs does not appear to be organized in a sex-dependent manner.

### Light-evoked vSUB aEPSC frequency is greater at D2R vs D1R MSN synapses

To further characterize this unique circuit, we next aimed to identify the underlying synaptic properties that contribute to the biased synaptic strength of vSUB-D2R MSN synapses. This bias can be a result of greater presynaptic release, greater postsynaptic strength, and/or more numerous synaptic contacts. First, we quantified paired pulse ratios (PPRs) of presynaptic vSUB boutons, an indirect measure of release probability, but found PPRs were not different between D1R and D2R MSNs (20 ms ISI: U = 96.3, p = 0.755, Mann-Whitney; 50 ms ISI: t(27) = 0.828, p = 0.415, unpaired t-test; 100 ms ISI: t(27) = 1.272, p = 0.214, unpaired t-test) (Figure 3A). Next, we assessed the quantal properties of these synapses by monitoring strontium-evoked asynchronous EPSCs (aEPSCs). We substituted extracellular calcium for strontium, which desynchronizes light-evoked vesicle release from vSUB terminals; resulting in a reduced peak EPSC followed by asynchronous release events that are quantal in nature (Bekkers & Clements, 1999). aEPSC amplitude is commonly thought to reflect postsynaptic strength, while aEPSC frequency may reflect presynaptic release probability and/or the number of active postsynaptic contacts (Sinnen et al., 2017). To assess if we could effectively discern vSUB-mediated aEPSCs from spontaneous activity in dorsal NAcMS MSN synapses, we compared baseline activity to aEPSCs occurring immediately after vSUB terminal stimulation and observed a 3-fold increase in aEPSC frequency (baseline vs light-evoked: t(11) = 8, p < 0.0001, paired t-test) (Figure 3B). While we did not observe differences in the amplitude of light-evoked vSUB aEPSCs, we did observe that aEPSC frequency was ∼2-fold higher at vSUB-D2R MSN synapses compared to D1R MSN synapses (amplitude: t(24) = 0.89, p = 0.383; frequency: t(24) = 7, p < 0.0001, unpaired t-tests; Cumulative frequency of amplitude and inter-event interval: D = 0.045 and 0.103, p = 0.2713 and p < 0.0001, respectively, K-S test) (Figure 3C). Taken together, the higher aEPSC frequency at vSUB-D2R MSN synapses cannot be explained by presynaptic release (Figure 3A) indicating that the biased evoked synaptic strength measured at vSUB-D2R MSN synapses (Figure 2C) is driven by an imbalance of synaptic contacts made by vSUB onto D2R MSNs and D1R MSNs (Figure 3C).

**Figure 3.**
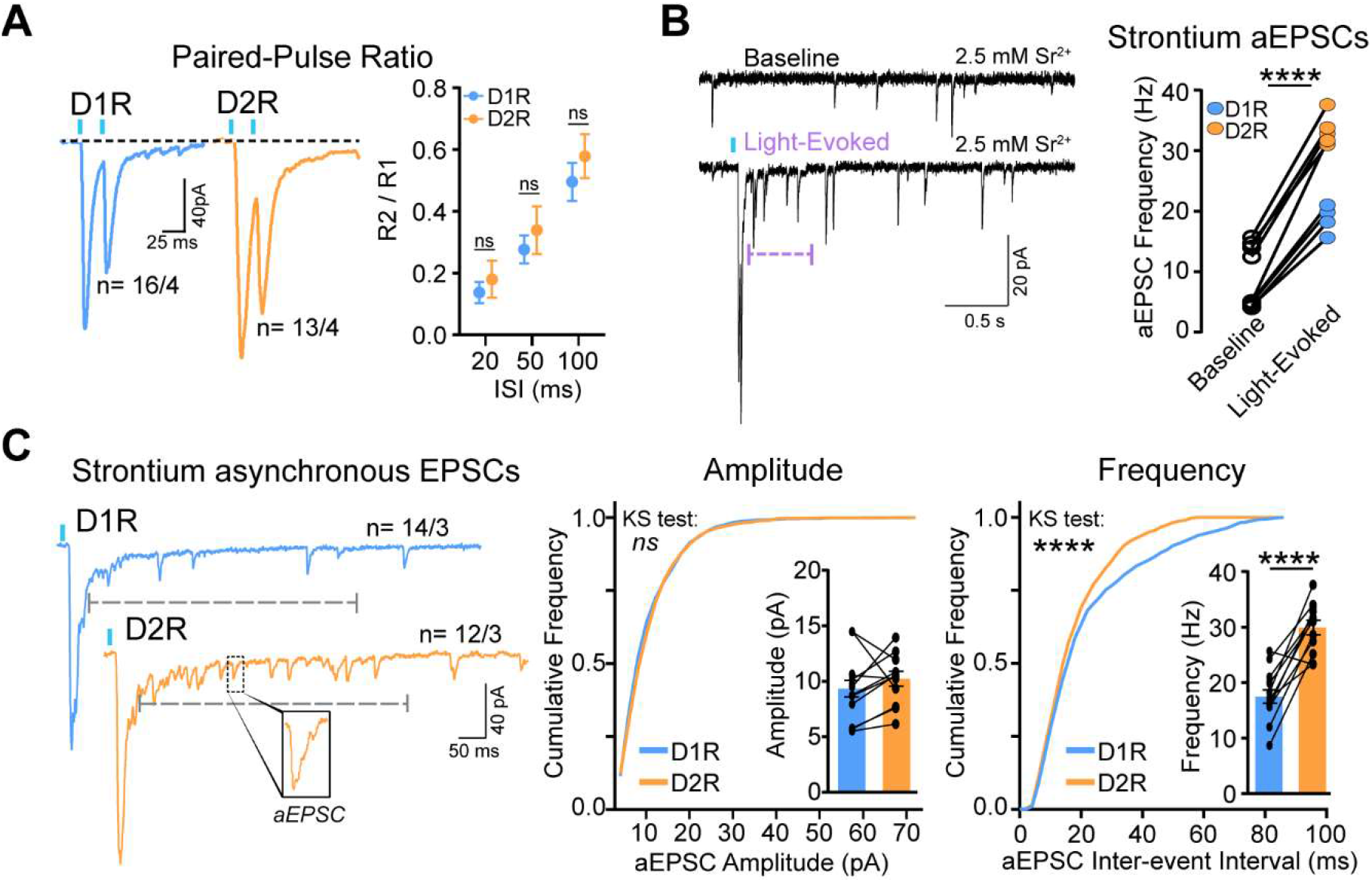
Light-evoked vSUB aEPSC frequency is greater at D2R vs D1R MSN synapses. (A) Presynaptic release probability is similar at vSUB-D1R and -D2R MSN synapses. Representative traces (left) and quantification (right) of light evoked PPRs from D1R and D2R MSNs. R2 / R1 (2nd response / 1st response). (B) Left: Representative traces of strontium-mediated asynchronous EPSCs from the same cell recorded in the absence of optical stimulation (Baseline; top) or after a single optical stimulation (Light-Evoked; bottom). Right: Summary plot of aEPSC frequency at baseline and after optical stimulation of vSUB synapses on D1R and D2R MSNs. aEPSC frequency of D2R MSNs increases significantly during 500 ms following light stimulation. n = 9 pairs. (C) Frequency but not amplitude of vSUB aEPSCs is greater in D2R compared to D1R MSNs. Left: Representative traces of strontium-mediated aEPSCs from a D1R and D2R MSN dual recording. Analysis of aEPSC amplitude and frequency was done in 500 ms window post stimulation (gray dashed line). See inset for a magnification of exemplary single aEPSC. Middle and Right: Cumulative frequency distribution of aEPSC amplitude and frequency, respectively, from D1R and D2R MSNs. Inset bar graphs show mean aEPSC amplitude and frequency. D1R and D2Rs from simultaneous dual recordings are connected. Error bars ± SEM. Number of cells and animals used for each experiment is included in each figure or corresponding figure legend.

### vSUB-NAcMS synapses harbor GluA2-lacking AMPA receptors

The unexpected bias of excitatory synaptic transmission onto D2R MSNs prompted us to further interrogate the synaptic properties of these inputs. We next assessed the properties of the receptors that populate vSUB-D1R and D2R MSN synapses, starting with the AMPAR subunit composition. We performed AMPAR current-voltage (I/V) experiments to examine the composition of AMPARs at these synapses. As a control, we first assayed total glutamatergic input onto MSNs by monitoring electrically evoked AMPAR EPSCs. Consistent with previous reports, we found that I/V curves were linear, indicating the majority of MSN synapses contain calcium-impermeable AMPARs (CI-AMPARs) (Figure 4A) (Britt et al., 2012; Pascoli et al., 2014; Terrier et al., 2016). Next, we assayed AMPAR composition at vSUB-MSN synapses by selectively activating vSUB terminals via optogenetic stimulation in drug-naïve animals. Surprisingly, AMPARs at vSUB-D1R and -D2R MSN synapses exhibited inward rectification at depolarized potentials in the presence of intracellular spermine, which is a hallmark of calcium-permeable AMPARs (CP-AMPARs) (Figure 4B). At glutamatergic synapses in NAc, CP-AMPARs are typically found following experience-dependent plasticity, and not in baseline conditions in drug-naïve animals. To confirm that the observed rectification was due to spermine-blockade of CP-AMPARs, we stimulated vSUB terminals and recorded from MSNs using an internal solution without spermine, and as expected, found that the resulting I/V curve and corresponding rectification index were significantly more linear (H = 10.61, p = 0.005, Kruskal-Wallis test; Dunn’s multiple comparisons: No spermine vs D1R: p = 0.009, No spermine vs D2R: p = 0.008, D1R vs D2R: p > 0.999) (Figure 4B). To further confirm that CP-AMPARs are expressed at these synapses, we washed-in the CP-AMPAR antagonist, NASPM, while optically stimulating vSUB terminals. Further supporting the idea that CP-AMPARs are basally present in drug-naïve vSUB-MSN synapses, we observed 34 ± 1% and 42 ± 3.5% depression of vSUB-AMPAR EPSC amplitudes in D1R and D2R MSNs, respectively, following 20 min of NASPM exposure (U = 7, p = 0.352, Mann-Whitney) (Figure 4C). The series resistances monitored throughout each experiment were stable (<2% deviation from baseline), indicating that the depression of light-evoked AMPAR-mediated currents was likely due to NASPM block of CP-AMPARS and not a result of cellular rundown or loss of voltage clamp. To our knowledge, the basal presence of CP-AMPARs at D1R and D2R MSN synapses in drug-naïve mice is novel and a unique property of vSUB output to NAcMS.

**Figure 4.**
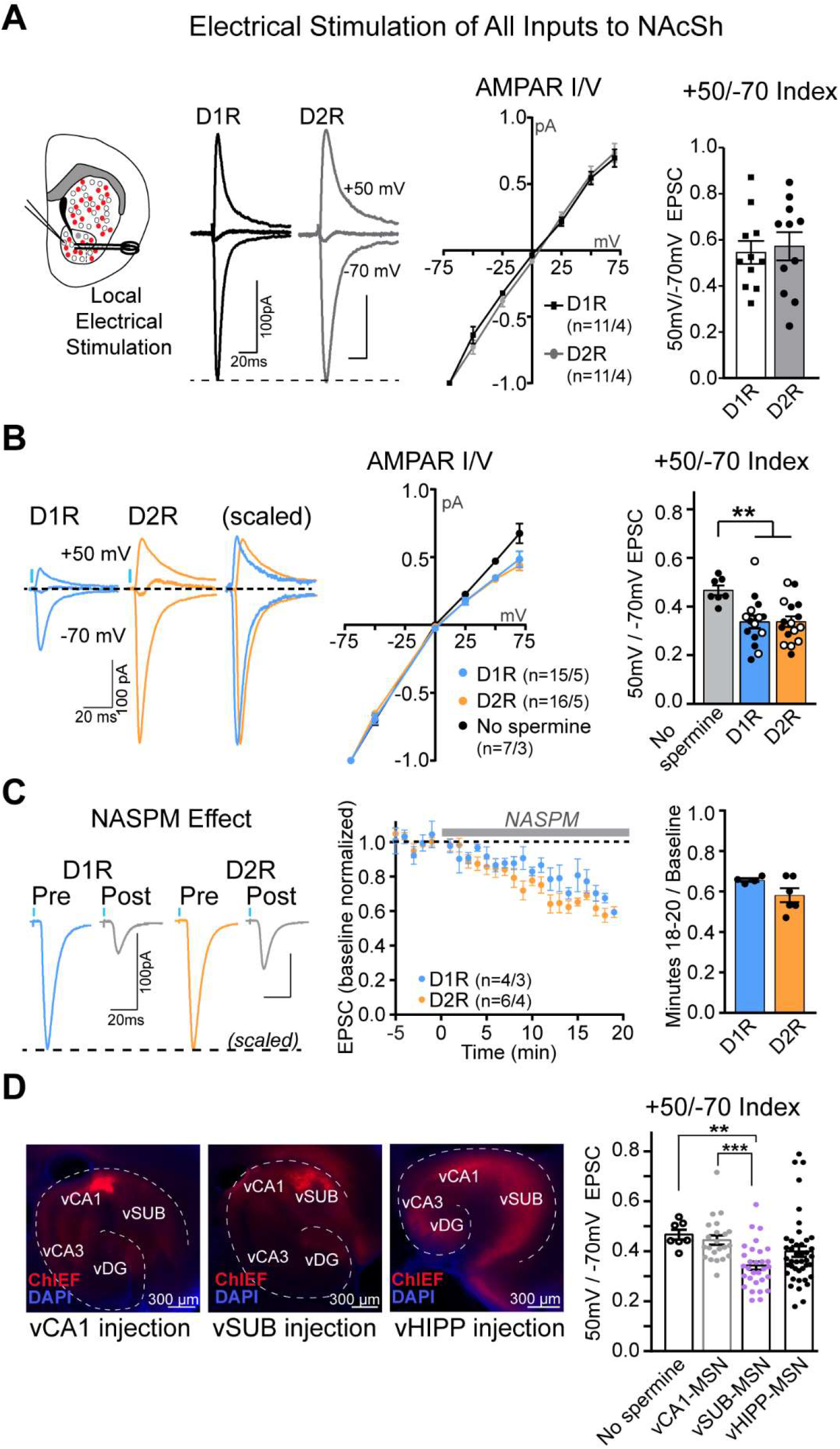
vSUB-NAc medial shell synapses harbor GluA2-lacking AMPA receptors. (A) General activation of glutamatergic synapses via local electrical stimulation reveals linear AMPAR current-voltage (I/V) relationships. From far left: Schematic drawing of experiment set-up, and representative traces of electrically evoked EPSCs voltage-clamped at −70, 0, and +50 mV (overlaid) from D1R and D2R MSNs. Middle: AMPAR I/V curves. Right: Summary graphs of the rectification indices for electrically evoked EPSCs from D1R and D2R MSNs. (B) AMPAR I/V relationships of vSUB-D1R and vSUB-D2R MSN synapses exhibit inward rectification. Left: Representative traces of AMPAR-mediated EPSCs at −70, 0, and 50 mV (overlaid) at vSUB to D1R and D2R MSN synapses. Middle and Right: AMPAR I/V curves and rectification indices (respectively) of MSNs patched without spermine in internal solution (black line), D1R MSNs (blue line), and D2R MSNs (orange line). There is no difference in D1R and D2R MSN rectification indices between females and males. (Females: closed circles, males: open circles) (C) Extracellular application of NASPM depresses vSUB-MSN AMPAR EPSC amplitudes. Left: Representative scaled traces of D1R and D2R MSN EPSCs before and after application of NASPM. Middle: EPSC amplitude normalized to 5-minute baseline and recorded following 18-20 minutes of NASPM application. Right: Summary graph of AMPAR EPSC amplitudes averaged over the last 2 minutes of NASPM wash-in reveals a ∼40% depression. (D) AMPAR rectification differs depending on presynaptic input. Left: Images of AAV-ChIEF-mRuby injections in vCA1 (left), vSUB (middle), and entire vHIPP (right). Scale bars: 300 μm. Right: Summary graph of rectification indices. No Spermine control: n=7 /3, vCA1: n=22 /3, vSUB: n=30 /5, vHIPP: n=42 /6. H = 21.69, p < 0.0001, Kruskal-Wallis test; Dunn’s multiple comparisons: vCA1 vs No spermine: p > 0.9999, vCA1 vs vSUB: p = 0.0005, vCA1 vs vHIPP: p = 0.0828, No spermine vs vSUB: p = 0.0035, No spermine vs vHIPP: p = 0.091, vSUB vs vHIPP: p = 0.351. Error bars ± SEM. Number of cells and animals used for each experiment is included in each figure or corresponding figure legend.

Our finding that vSUB-NAcMS MSN synapses contain CP-AMPARs is particularly unexpected because optical stimulation of fibers from *total* vHIPP identified that these synapses contain CI-AMPARs (Britt et al., 2012; Pascoli et al., 2014). However, vHIPP output to NAcSh is a composite of vCA1 and vSUB, thus the presence of CI-AMPARs observed at vHIPP-MSN synapses might primarily reflect input originating from vCA1. To attempt to reconcile the differences in AMPAR composition at vSUB- and vHIPP-MSN synapses, we selectively injected vCA1 with AAV-ChIEF (Figure 4D) and assessed the AMPAR I/V relationship. We found that the AMPAR rectification index at vCA1-MSN synapses was significantly higher than that of the inwardly rectifying vSUB-MSNs, and was indistinguishable from the no-spermine control, suggesting that vCA1 synapses are populated by CI-AMPARs (Figure 4D). Furthermore, we performed *total* vHIPP injections with AAV-ChIEF, performed I/V experiments in MSNs. We validated that all subfields of vHIPP were infected with AAV-ChIEF and found that the vHIPP rectification index fell between the vCA1 and vSUB rectification indices (RI: vCA1 = 0.45 ± 0.02, vSUB = 0.34 ± 0.02, vHIPP = 0.40 ± 0.02) and had the largest spread of the three conditions (coefficient of variation: vCA1 = 20%, vSUB = 26%, vHIPP = 35%) (Figure 4D). Therefore, the AMPAR composition monitored at vHIPP-MSN synapses resembles a mixture of vCA1 and vSUB input. Together, the I/V relationship and NASPM experiments strongly suggest that in drug-naïve animals, vSUB-MSN synapses contain CP-AMPARs, and that this property is unique from vCA1-MSN synapses (H = 21.69, p < 0.0001, Kruskal-Wallis; Dunn’s multiple comparisons: vCA1 vs vSUB: p = 0.0005, No spermine vs vSUB: p = 0.0035) (Figure 4D).

Additionally, the comparison of AMPAR rectification indices at vCA1, vHIPP with vSUB synapses suggests that differences in injection site (e.g., predominant infection of vCA1) likely results in the disproportionate representation of vCA1-MSN synaptic properties and provides an explanation why CI-AMPARs have been observed at vHIPP-NAcMS MSN synapses. This unique feature of vSUB synapses prompted us to ask if this phenotype is sexually dimorphic. We separated our I/V data by sex and found both males and females exhibit the same degree of inward rectification at vSUB-D1R and -D2R synapses (effect of sex: ns, effect of cell-type: ns, effect of interaction: ns, Ordinary Two-way ANOVA) (Figure 4B, females: closed circles, males: open circles).

### NMDA receptor function is equal at vSUB-D1R and -D2R MSN synapses

Next, we interrogated the properties of NMDA receptors (NMDARs) at vSUB-NAcMS synapses. NMDARs are required for the induction of synaptic plasticity at vHIPP-NAcMS synapses following cocaine exposure or high frequency stimulation (MacAskill et al., 2014; Pascoli et al., 2014; LeGates et al., 2018). Interestingly, at vHIPP synapses made on unidentified MSNs, NMDARs are less sensitive to blockade by Mg^2+^ and exhibit strong inward current at negative holding potentials compared to inputs from basolateral amygdala or medial prefrontal cortex, indicating that NMDAR subunit composition is input-specific (Britt et al., 2012). Given that CP-AMPARs are unique to vSUB-MSN synapses, we questioned if vSUB synapses in NAcMS also contain NMDARs with strong inward current and whether NMDAR composition exhibits synapse-specificity. To specifically address these questions, we pharmacologically isolated NMDAR-mediated EPSCs while optically stimulating vSUB terminals. First, we assessed the NMDAR I/O relationship at vSUB-MSN synapses and found that amplitudes and the corresponding I/O curves were similar between D1R and D2R MSNs (U = 89, p = 0.7168, Mann-Whitney) (Figure 5A). Next, we assayed NMDAR subunit composition by testing the I/V relationship. D1R and D2R MSNs NMDAR I/V curves overlapped and displayed very minimal inward current at negative potentials (fraction of +50 mV amplitude: 0.14 at −70 mV and 0.25 at −25 mV) (Figure 5B). Thus, vSUB-D1R and -D2R MSN synapses contain more canonical NMDARs compared to *total* vHIPP inputs suggesting that vSUB inputs may be less excitable, and that the induction of plasticity at vSUB-NAcMS synapses could be distinct from vHIPP-NAcSh synapses. Together, our results indicate that both AMPAR- and NMDAR-mediated synaptic transmission properties at vSUB inputs to D1R and D2R MSNs are distinct from vHIPP-MSN synapses.

**Figure 5.**
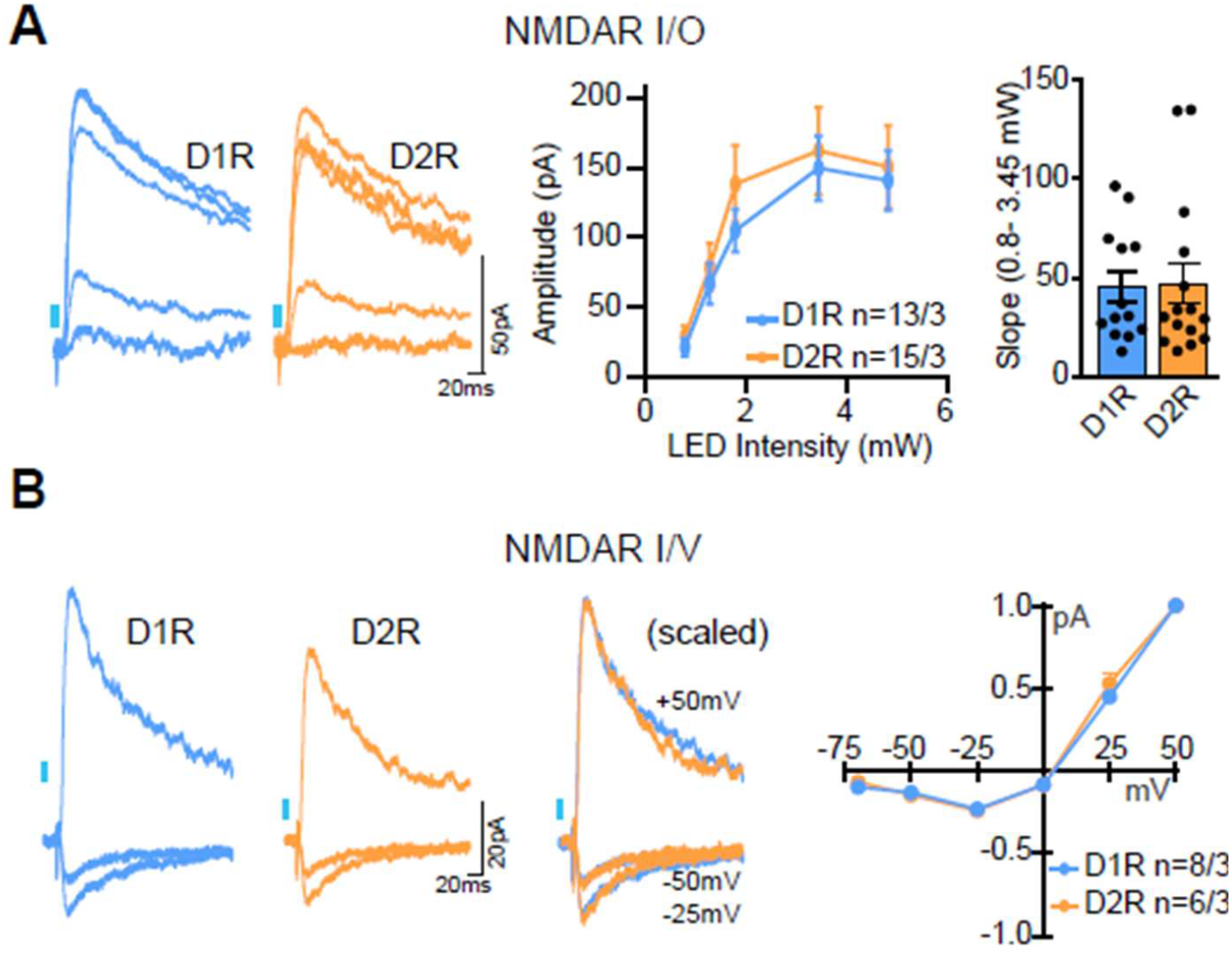
NMDA receptor function is equal at vSUB-D1R and -D2R MSN synapses. (A) Optically evoked NMDAR-mediated EPSC amplitudes are equal at vSUB to D1R and D2R MSN synapses. Left: MSNs are voltage-clamped at +50 mV, and EPSCs are recorded at different light intensities. D1R/D2R representative traces are overlaid. Middle: NMDAR-mediated I/O curve. Right: Summary graph of NMDAR I/O slopes. (B) NMDAR I/V relationships. Left: Representative NMDAR-mediated EPSCs at −25, -50, and +50 mV. Right: NMDAR I/V curves at vSUB-D1R and D2R MSNs. All values are normalized to +50 mV. Error bars ± SEM. Number of cells and animals used for each experiment is included in each figure.

### Ventral subiculum to D2R MSN synapses are potentiated following cocaine administration

vHIPP-NAc glutamatergic inputs undergo plasticity following cocaine administration and withdrawal. In drug-naïve mice, vHIPP exhibits a bias in synaptic strength for D1R MSNs and vHIPP-D1R synapses are selectively potentiated following withdrawal from contingent and non-contingent administration of cocaine. However, whether vSUB-MSN synapses undergo similar drug-induced plasticity is untested. Our results thus far indicate that vSUB possess distinct basal synaptic transmission properties relative to *total* vHIPP or vCA1 at MSN synapses. We therefore tested the impact of cocaine exposure on vSUB-MSN synapses. We injected D1R-TdTomato mice with AAV-ChIEF on ∼P21 then administered daily intraperitoneal injections of either cocaine (20 mg/kg) or saline for 5 days. After a 10–11 day withdrawal period, we interrogated the synaptic properties of vSUB-D1R and vSUB-D2R synapses in dorsal NAcMS *ex vivo* slices (Figure 6A-B). We systematically tested for the manifestation of cocaine-induced pre- and post-synaptic plasticity by measuring PPRs to assess changes in presynaptic release, AMPA-NMDA ratios to assess postsynaptically expressed potentiation, and AMPAR I/V curves and NASPM wash-ins to detect changes in the subunit stoichiometry of AMPARs. Presynaptic release was unaffected by cocaine withdrawal, as PPRs were comparable between saline or cocaine injected animals at vSUB-D1R and vSUB-D2R synapses (D1R saline vs D1R cocaine: t(60) = 1.5, p = 0.150, D2R saline vs D2R cocaine: t(64) = 0.15, p = 0.882, unpaired t-tests) (Figure 6C). Next, we measured AMPA-NMDA ratios by quantifying the EPSC amplitudes at holding currents of −70 mV and +40 mV to isolate AMPAR- and NMDAR-mediated EPSC amplitudes, respectively. Interestingly, while we did not observe cocaine-induced changes in the A/N ratio at vSUB-D1R synapses, we observed a significant increase in the A/N ratio at vSUB-D2R synapses in cocaine injected animals (D1R saline vs D1R cocaine: t(55) = 0.74, p = 0.462, D2R saline vs D2R cocaine: t(52) = 2.314, p = 0.025, unpaired t-tests) (Figure 6D). The synaptic potentiation observed at vSUB-D2R synapses represents another unique property of these synapses and contrasts with the selective cocaine-induced potentiation at vHIPP-D1R synapses. We next tested if the cocaine-induced plasticity changes the composition of AMPARs at vSUB-D2R synapses. We found that the I/V rectification indices at either D1R or D2R synapses was unchanged between cocaine compared to saline-injected groups (D1R saline vs D1R cocaine: t(30) = 0.297, p = 0.769, D2R saline vs D2R cocaine: t(32) = 0.559, p = 0.580, unpaired t-tests) (Figure 6E-G). We confirmed this finding with application of extracellular NASPM, which produced a similar degree of AMPAR-mediated EPSC depression in saline and cocaine injected animals at both vSUB-D1R and -D2R synapses (D1R saline vs D1R cocaine: U = 10, p = 0.43, D2R saline vs D2R cocaine: U = 9, p = 0.329, Mann-Whitney) (Figure 6H-J). Together, these results indicate that while cocaine-induced plasticity at vSUB-D2R synapses manifests as an increase in synaptic AMPARs, the stoichiometry of CI- and CP-AMPARs established basally is unexpectedly maintained after synaptic potentiation.

**Figure 6.**
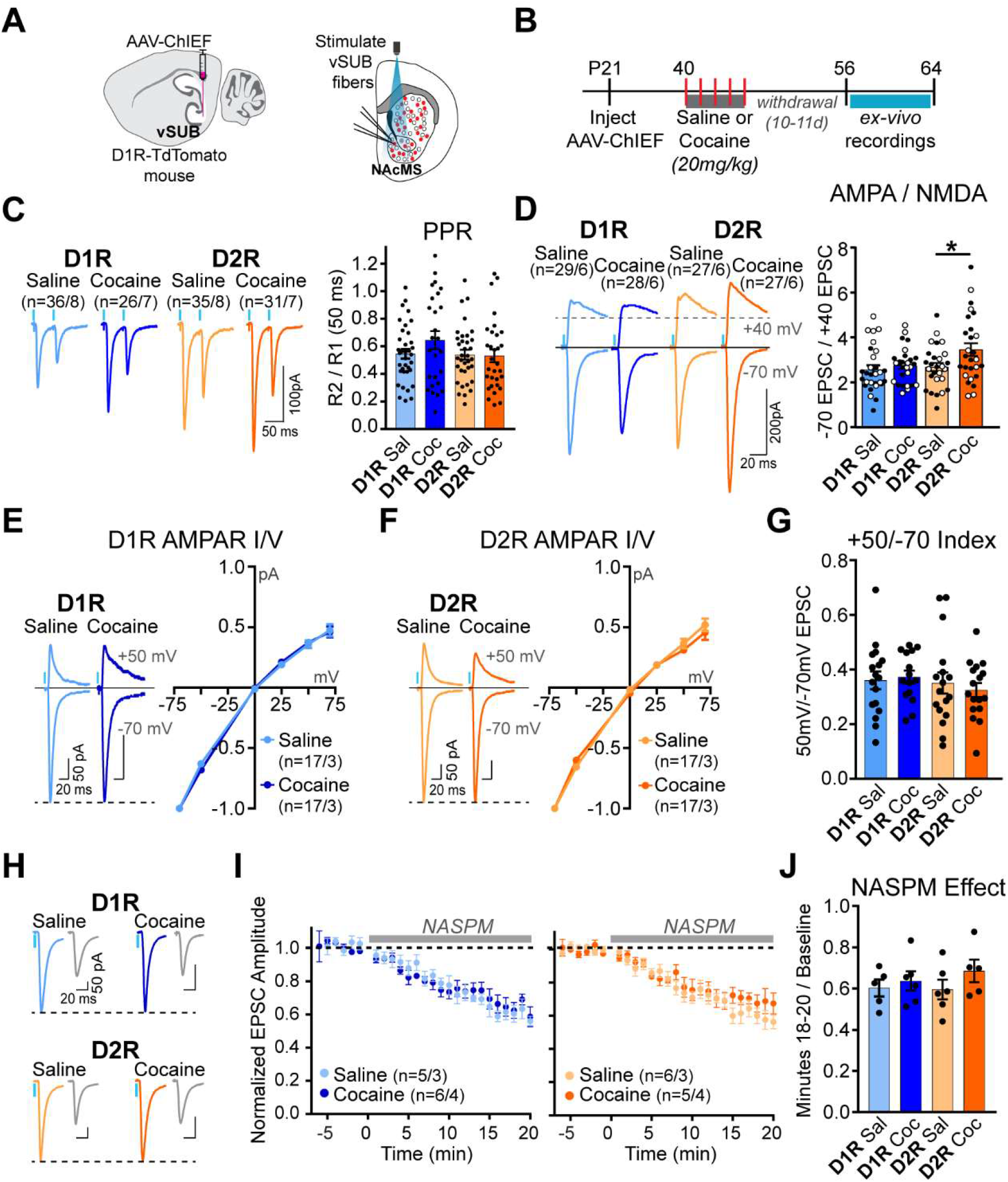
Cocaine potentiates vSUB-D2R MSN synapses. (A) Simultaneous dual recording schematic to monitor vSUB-D1R and -D2R MSN synaptic properties. AAV-ChIEF was injected in vSUB of D1R-TdTomato mice, then vSUB input to D1R and D2R MSNs in NAcMS was measured by optically stimulating vSUB terminals. (B) Experimental timeline: Mice were injected with AAVs on P21 then 3 weeks later received daily non-contingent injections of saline or cocaine (20 mg/kg) or saline for 5 days. Animals were sacrificed for *ex vivo* recordings following a 10 to 11-day withdrawal period. (C) Exposure to cocaine does not change release probability in vSUB-NAcMS synapses. Left: Representative traces of light-evoked paired-pulse ratios (PPRs) from D1R and D2R MSNs in cocaine and saline injected mice. Right: Quantification of PPRs. R2 / R1 (2nd response / 1st response). 50 ms inter-stimulus interval. (D) Exposure to cocaine induces synaptic plasticity at vSUB-D2R MSN synapses. Left: Representative traces of light evoked EPSCs at −70 and +40 mV from D1R and D2R MSNs in cocaine and saline injected mice. Right: Quantification of AMPA/NMDA ratios. A direct sex-specific comparison of AMPA/NMDA ratios did not identify sex differences in cocaine-induced plasticity (females: closed circles, males: open circles). Sample size: Female N = 3 mice, D1R saline n=14, D1R cocaine n=15, D2R saline n=15, D2R cocaine n=16. Male N = 3 mice, D1R saline n=15, D1R cocaine n=13, D2R saline n=12, D2R cocaine n=11. (E and F) Left: Representative traces of AMPAR EPSCs at −70 and +50 mV. Right: AMPAR I/V plots of saline and cocaine injected mice from vSUB-D1R and -D2R MSN synapses, respectively. (G) Exposure to cocaine does not change AMPAR inward rectification compared to saline controls. Rectification index of D1R and D2R MSNs in saline and cocaine injected mice. (H) Representative traces of AMPAR EPSCs before and after application of NASPM. (I) Change in EPSC amplitude normalized to baseline over time in D1R (left) and D2R MSNs (right) from saline and cocaine injected mice. (J) Summary graph of EPSC amplitudes averaged over the last 2 minutes of NASPM wash-in from D1R and D2R MSNs in saline and cocaine injected mice. Error bars ± SEM. Number of cells and animals used for each experiment is included in each figure or corresponding figure legend.

Finally, we tested if the vSUB phenotype was sex-specific by separating our A/N data by sex and performing a three-way ANOVA (Figure 6D, females: closed circles, males: open circles). As expected, there were main-effects of cell-type and drug condition, but there was no main-effect of sex, nor any interactions between the three variables (Main effects: sex: p = 0.16, cell-type: p = 0.0339, drug: p = 0.0209; interactions: sex-cell: p = 0.74, sex-drug: p = 0.14, cell-drug: p = 0.16, sex-cell-drug: p = 0.29, Three-way ANOVA). Thus, consistent with our findings that the vSUB-NAcMS circuit is unique from the *total* vHIPP to NAcSh circuit, we find that vSUB inputs to dorsal NAcMS display unique cocaine-induced plasticity distinct from *total* vHIPP input (which includes vSUB but also vCA1 and possibly vEC), which experiences potentiation at D1R synapses, whereas vSUB-specific input undergoes D2R MSN specific alterations.

## Discussion

vHIPP input to the NAcMS has received attention for its proposed role in SUDs. vCA1 and vSUB are surprisingly distinct subregions in vHIPP – they differ in principal neuron diversity, projection targets within NAcSh, and in their influence on behavior – however, a distinction between their functional contribution in the vHIPP-NAcMS circuit has not been explored. Here, we systematically interrogated the cell-type-specific organization and synapse-specific properties of the vSUB-NAcMS circuit and report four fundamentally important findings. First, using an intersectional retrograde circuit tracing approach, we defined the connectivity of RS and BS neurons with D1R and D2R MSNs and found that equal proportions of RS and BS neurons innervate D1R and D2R MSNs. Second, we performed the first functional dissection of vSUB-NAcMS circuitry and found a striking bias of excitatory synaptic strength at D2R synapses, relative to D1R synapses, mediated by differences in synapse density. Third, we reveal that vSUB-D1R and -D2R synapses basally contain CP-AMPARs. Fourth, we report that vSUB-D2R synapses, but not D1R synapses, are selectively potentiated following cocaine administration. Importantly, the functional properties of vSUB-D1R and -D2R synapses in drug-naïve and drug exposed mice reveal novel properties of these synapses not previously observed in vHIPP-NAc synapses.

### Cell-type-specific organization and synaptic properties of the vSUB-NAcMS circuit

RS neurons represent about half of the total population of principal neurons in vSUB, however, RS cells provide ∼70% of vSUB output to total NAc. We used intersectional retrograde tracing to reveal that equal proportions of RS and BS neurons project to D1R and D2R MSNs. Our findings suggest that RS afferents may primarily innervate non-MSNs, such as local interneurons that provide strong inhibition within NAcMS (Baimel et al., 2022; Scudder et al., 2018). Although the functional relevance of RS and BS neurons is largely unknown, RS and BS neurons exhibit distinct forms of long-term potentiation and the activation of D1/D5 receptors on RS neurons reduces the threshold for LTP induction at CA1-SUB synapses (Wozny et al., 2008; Roggenhofer et al., 2013). Additionally, the intrinsic excitability properties of RS neurons undergo experience-dependent plasticity following exposure to novel contexts relevant for contextual fear learning (Dunn et al., 2018). Thus, RS neuron input to local interneurons in NAcMS may serve as an important feed-forward mechanism to sustain excitatory-inhibitory balance of MSNs following cocaine exposure. Future experiments are needed to elucidate the functional properties of RS input onto NAcMS local interneurons.

We also report that vSUB neurons exhibit a synaptic bias for D2R MSNs in dorsal NAcMS. Furthermore, vSUB-D1R and -D2R synapses unexpectedly contain CP-AMPARs, which to our knowledge, is a unique property for synapses in NAc. By contrast, previous studies of vHIPP-MSN synapses revealed a bias for D1R MSNs and the presence of synaptic CI-AMPARs (Britt et al., 2012; MacAskill et al., 2014; Pascoli et al., 2014). To reconcile the striking differences in AMPAR composition, we found that CI-AMPARs populate vCA1-MSNs synapses suggesting that the previously reported synaptic properties of vHIPP-NAc synapses may primarily reflect the properties of vCA1 synapses. vCA1 and vSUB are molecularly and functionally distinct subregions of vHIPP. vCA1 neurons encode reward location, cocaine CPP, and store social memory (Ciocchi et al., 2015; Okuyama et al., 2016; Zhou et al., 2019) while vSUB RS and BS principal neurons encode the vigor of reward seeking (Britt et al., 2012; Lindenbach et al., 2022; Pascoli et al., 2014). Perhaps it is not surprising that these regions exhibit differences in basal synaptic transmission properties. The striking differences in CI- and CP-AMPAR usage at vCA1 and vSUB synapses in NAcMS highlights the importance of studying each region separately.

### Functional implications of the synaptic properties unique to vSUB

In NAc, CP-AMPARs typically manifest after drug exposure, thus, the use of CP-AMPARs at vSUB-MSN synapses of drug-naïve mice is particularly intriguing and raises the question – what purpose does the basal expression of these CP-AMPARs serve? The basal expression of CP-AMPARs may maintain vSUB-MSN synapses in a metaplastic state to facilitate future plasticity (Park et al., 2021). Additionally, it has been proposed that synaptic plasticity can be triggered by Ca^2+^ influx via CP-AMPARs, independent of postsynaptic NMDARs (Lamsa et al., 2007; Mameli et al., 2011). This form of plasticity requires presynaptic excitatory transmission paired with postsynaptic hyperpolarization. Thus, feed-forward local inhibition in NAc, perhaps driven by vSUB RS neurons, may be required to induce synaptic plasticity at vSUB-MSN synapses. While CP-AMPARs alone can promote LTP, Ca^2+^ influx through CP-AMPARs and NMDARs at the same synapse can additively induce LTP, which suggests that vSUB-MSN synapses can undergo exaggerated LTP in response to relevant stimuli (Jia et al., 1996). By contrast, vHIPP-MSN synapses contain NMDARs that are less sensitive to Mg^2+^ block, and display significant inward current at hyperpolarized potentials (Britt et al., 2012). The NMDARs at vHIPP synapses may function in a similar manner to the CP-AMPARs at vSUB synapses to facilitate the induction of plasticity.

### Synapse-specific cocaine induced plasticity

We observed a selective cocaine-induced potentiation of vSUB-D2R synapses. By contrast, cocaine plasticity occurs selectively at vHIPP-D1R synapses (Britt et al., 2012; MacAskill et al., 2014; Pascoli et al., 2014). It is commonly thought that D1R and D2R MSNs participate in opposing but parallel pathways to regulate drug seeking behavior – D1R plasticity strictly promotes reward behavior whereas plasticity at D2R synapses encodes aversion. Thus, based on these putative roles, our findings seemingly contradict with the notion that hyperactivity of vSUB promotes drug seeking and drug reinstatement. However, our results add to a growing literature that NAc D2R MSNs play more complex roles in drug-induced plasticity and motivated behavior than previously appreciated and, in many cases, promote reward seeking behavior (Terrier et al., 2016; Inbar et al., 2022; Sjulson et al., 2018; Soares-Cunha et al., 2016; Gong et al., 2021; Cole et al., 2018). While NAc D2R MSNs mostly project to inhibitory neurons in VP to encode aversive behavior, D2R MSNs in dorsal NAcMS promote reward via specific projections to excitatory neurons in VP (Yao et al., 2021). vSUB robustly projects to dorsal NAcMS, which is where we exclusively performed our experiments. Thus, vSUB may preferentially control the basal activity and drug-induced plasticity of a specific subset of D2R MSNs that promote reward seeking behavior. The selective potentiation of vSUB-D2R MSN synapses has not been described for vHIPP input and further underscores the need to study each output region of vHIPP individually.

In most subregions of striatum, dopamine signaling, acting via D1-receptors, is required for and enhances LTP on D1R-expressing MSNs, whereas activation of D2-receptors may inhibit LTP (Goto & Grace, 2005; Håkansson et al., 2006; Iino et al., 2020; LeGates et al., 2018; Pawlak & Kerr, 2008; Schotanus & Chergui, 2008). Synaptic D1- and D2-receptors modulate excitability via G-protein signaling and can form heterodimers with the NMDAR subunit NR2B to either facilitate or inhibit NMDAR function (X.-Y. Liu et al., 2006; Pascoli et al., 2014; Cahill et al., 2014). This raises the question: if dopamine signaling is required for LTP in NAc, how can we observe selective LTP at vSUB-D2R MSN synapses? However, at vHIPP-NAcMS synapses, D1- and D2-receptor activation are neither required for nor do they enhance MSN LTP (LeGates et al., 2018). Dopamine signaling-independent LTP may be due to the fact that in NAcMS, dopamine terminals mostly contact MSN dendrites instead of spines, suggesting that dopamine signaling may be less able to directly influence glutamatergic input to spines (Zahm, 1992; Meredith et al., 2008). If LTP in dorsal NAcMS is independent of dopamine signaling, how do vSUB-D2R synapses selectively potentiate after cocaine exposure? It is possible that only vSUB neurons projecting to D2R MSNs experience increased activity following cocaine exposure, resulting in downstream potentiation. Additionally, microcircuit differences within NAc may drive synapse-specific changes. Biased feedforward inhibition onto D2R MSNs in dorsal NAcMS could provide the hyperpolarization necessary to enable potentiation at synapses that contain CP-AMPARs. Alternatively, vSUB-D2R synapses could be molecularly unique in other, yet untested ways. For example, cell adhesion molecules can specify synapse identity and govern synaptic plasticity (Südhof, 2017) and although largely unstudied in NAc, cell-type-specific expression and/or localization of cell adhesion molecules may facilitate LTP at D2R-expressing MSN synapses (Fuccillo et al., 2015; Rothwell et al., 2014).

### vSUB-D1R and -D2R synapses are not sexually dimorphic

Our recent work identified sex differences in PV-mediated inhibition of vSUB RS vs BS neurons and here we tested whether vSUB synapses in dorsal NAcMS also exhibit sexual dimorphism (Boxer et al., 2021). We assessed sex differences throughout our study, but found no differences in any of the circuit or synaptic properties measured. Importantly, these measurements were performed using terminal stimulation, which drives presynaptic release at vSUB terminals independent of real-time regulation by the vSUB local circuit. Sex specific differences are reported for SUDs in humans and reward seeking behaviors in rodents (Becker, 2016), however, a comprehensive understanding whether the sex-specific properties of vSUB local inhibition contribute to sexually dimorphic reward seeking behaviors is unknown. Future studies into the intact vSUB-NAcMS circuitry conducted in behaving animals will be critical to determine how vSUB local circuitry regulates excitatory output to NAcMS.

## Methods

### Animals

Male and female mice were bred at the University of Colorado Anschutz and were from a B6;129 or B6.Cg mixed genetic background. The B6.Cg-Tg(Drd1a-tdTomato)6Calak/J (“D1R-tdTomato,” Jax 016214) breeders, B6;129-Tg(Drd1-cre)120Mxu/Mmjax (“D1R-Cre,” #037156-JAX) breeders, and the B6.FVB(Cg)-Tg(Adora2a-cre)KG139Gsat/Mmucd (“A2a-Cre,” MMRRC 036158-UCD) breeders were generous gifts from Dr. Robert Malenka. Mice were genotyped in house. Mice were housed in a dedicated animal care facility maintained at 35% humidity, 21-23°C, on a 14/10 light/dark cycle and housed in groups of 2-5 in ventilated cages with same-sex littermates with food and water *ad libitum*. Mice were stereotactically injected on P21-24, and electrophysiology experiments were performed at P56-65 in visibly healthy animals. For experiments related to. All experiments were carried out in cohorts, wherein at least two littermates were injected with either saline or cocaine. At P40-45, mice received intraperitoneal injections of cocaine (20 mg/kg) or saline in the home cage for five consecutive days. All procedures were conducted in accordance with guidelines approved by Administrative Panel on Laboratory Animal Care at University of Colorado, Anschutz School of Medicine, accredited by Association for Assessment and Accreditation of Laboratory Animal Care International (AAALAC) (00235).

### Viral constructs

AAV vectors were constructed from an empty AAV transfer plasmid where the expression cassette is as follows: left-ITR, human synapsin promoter, multiple cloning site, WPRE and right ITR. Cre-dependent plasmids contained: left-ITR, human synapsin promoter, 5’ LoxP site, multiple cloning site, 3’ LoxP site, WPRE and right ITR. Plasmids carrying mRuby and T2A cDNAs were generous gifts from Dr. Kevin Beier (Beier et al., 2017). To generate AAVs, HEK293T cells were transfected with a AAV transfer plasmid, pHelper and pRC-DJ or pRC-retro. AAVs were purified as previously described (Aoto et al., 2013). Briefly, 72-hour post transfection, cells were harvested, lysed and virus was purified and harvested from the 40% iodixanol fraction after ultracentrifugation. Virus was concentrated in a 100K MWCO Amicon filter. The following AAVs were used: AAV_DJ_-hSYN-ChIEF-mRuby; AAV_DJ_-hSYN-mCherry-T2A-WGA:Cre; AAV_DJ_-hSYN-DIO^LoxP^-mRuby; AAV_DJ_-hSYN-DIO^LoxP^-WGA:Flp-T2A-GFP; AAV_DJ_-hSYN-DIO^Frt^-mRuby; AAV_2_-Retro-mRuby

### Stereotactic surgeries

Stereotactic injections were performed on P21-24 mice. Animals were anesthetized with an intraperitoneal injection of 2,2,2-Tribromoethanol (250 mg/kg) then head fixed to a stereotactic frame (KOPF). Solutions containing AAVs (0.2-0.5 μL) were injected into ventral subiculum, ventral CA1, or nucleus accumbens medial shell at a rate of 10 μL/hr using a syringe pump (World Precision Instruments). Coordinates (in mm) were vSUB: anterior-posterior: −3.4, mediolateral: ±3.2 (relative to Bregma), and dorsoventral: −3.45 (relative to pia); vCA1: AP: −3.1, ML: ±3.2, DV: −3.45; NAcMS: AP: 1.5, ML: ±0.75, DV: −4.0.

### *Ex vivo* whole-cell electrophysiology

At P56-65, animals were deeply anesthetized with isoflurane and decapitated. Brains were rapidly dissected and 300 μm horizontal slices (for vSUB recordings) or coronal slices (for NAc recordings) were sectioned with a vibratome in cutting solution then moved to ACSF, as previously described (Boxer et al., 2021), were superfused with 29.5°C oxygenated ACSF containing (in mM) 126 NaCl, 26.2 NaHCO_3_, 11 D-Glucose, 2.5 KCl, 2.5 CaCl_2_, 1.3 MgSO_4_-7H_2_O, and 1 NaH_2_PO_4_. For NAc recordings, ACSF included 100 μM picrotoxin. To isolate AMPAR-mediated currents (AMPA I/O, strontium, and I/V experiments) 50 μM D-AP5 was also included in the ACSF, whereas NMDAR-mediated currents (NMDA I/O and I/V experiments) were isolated with 10 μM NBQX. For strontium experiments, CaCl_2_ was replaced with 2.5 mM SrCl_2_ in the recording ACSF. 20 μM NASPM was used for wash-in experiments.

MSNs in NAc were identified by their shape and size, high membrane resistance (>800 mΩ), and determined as D1R^+^ or D1R^−^ (putative D2R) by expression of tdTomato. Cells were visualized using an Olympus BX51W microscope and with a 40x dipping objective collected on a Hamamatsu ORCA-Flash 4.0 V3 digital camera using an IR bandpass filter.

Neurons were voltage-clamped at −70 mV in whole-cell configuration with a cesium-based internal solution containing (in mM) 117 cs-methanesulfonate, 15 CsCl, 10 TEA-Cl, 10 HEPES, 10 Phosphocreatine, 8 NaCl, 4 Mg_2_-ATP, 1 MgCl_2_, 0.5 Na_2_-GTP, and 0.2 EGTA. For A/N ratio, cells were held at −70 mV or +40 mV from the experimentally determined reversal potential to record AMPAR-mediated currents and NMDAR-mediated currents, respectively. 10 μM spermine was added to the internal solution for AMPA I/V experiments; 5 mM QX-314 was added for A/N Ratio and I/V experiments. To optogenetically stimulate ChIEF expressing vSUB fibers, slices were illuminated with 470 nm LED light (ThorLabs M470L2-C1) for 3 ms through the 40x dipping objective located directly over the recorded cell. With an illumination area of 33.18 mm^2^ the tissue was excited with an irradiance of 0.006 to 0.17 mW/mm^2^. To electrically stimulate MSN synapses, a homemade nichrome stimulating electrode was placed ∼200 μm from the patched cell and pulsed at 0.1 Hz at 100 µA (A-M Systems 2100 Isolated pulse stimulator).

For RS/BS ratio in vSUB, the ACSF contained no drugs, and mRuby^+^ neurons were patched in whole-cell configuration with a K-gluconate based internal containing (in mM): 95 K-gluconate, 50 KCl, 10 HEPES, 10 Phosphocreatine, 4 Mg_2_-ATP, 0.5 Na_2_-GTP, and 0.2 EGTA. vSUB pyramidal neuron identity (RS vs BS) was determined by the suprathreshold action potential firing pattern, as previously described (Boxer et al., 2021). All recordings were acquired using Molecular Devices Multiclamp 700B amplifier and Digidata 1440 digitizer with Axon pClamp™ 9.0 Clampex software, lowpass filtered at 2 kHz and digitized at 10-20 kHz.

### Analysis of electrophysiology recordings

Evoked EPSC peak amplitudes were determined using Axon™ pClamp10 Clampfit software. 12-20 sweeps (0.1 Hz) were averaged to obtain peak amplitude at each light intensity (I/O) or voltage (I/V). Input/output slope was calculated using the SLOPE function in Microsoft Excel: (pA amplitude range/mW intensity range). Release probability was assessed by measurements of paired-pulse ratios (PPRs) at inter-stimulus intervals of 20-100 ms. PPR was measured by dividing the average EPSC amplitude evoked by the second stimulus, by the average amplitude evoked by the first stimulus (R2/R1). A/N ratio was determined by dividing the peak EPSC amplitude at −70 mV (AMPA), by the EPSC amplitude measured at 50 ms post onset, at +40 mV (NMDA). Asynchronous EPSC event amplitudes (strontium experiments) were analyzed using Clampfit event detection software.

### Experimental Design and Statistical Analyses

Both female and male mice were used in this study. In relevant graphs, data from females are represented as closed circles while those from males are represented as open circles. Number of animals (N) and cells (n) used for each experiment is included in each figure or corresponding figure legend. All experiments were replicated in at least 3 animals. Experimenter was not blinded to animal sex, genotype, or cell identity, but was blinded to saline or cocaine assignments during data acquisition. No statistical method was used to predetermine sample size prior to the study. All data was tested for normality using D’Agostino & Pearson normality tests. If datasets exhibited normal distribution, Student’s paired and unpaired t tests and Two-way and Three-way ANOVAs were used to determine statistical differences. Otherwise, Mann-Whitney or Kruskal-Wallis tests were used. Tests were corrected for multiple comparisons using Sidak’s or Dunn’s multiple comparisons tests. For strontium datasets, Kolmogorov-Smirnoff tests were used to compare the cumulative probability plots. Differences were considered statistically significant when p< 0.05. Unless otherwise stated, all bar graphs are presented as mean ± SEM. Statistical tests and graph making were performed using Prism 7 (GraphPad), and figures were compiled in Adobe Illustrator.

## Acknowledgements

We would like thank the members of the Aoto lab for helpful discussions. We thank Nora Langer and Melanie Becher for their contributions to mice husbandry and AAV injections. We also thank Dr. Kevin Beier for contributing plasmids and Dr. Robert Malenka for contributing mouse lines. This work was supported by grants from the NIH: R00MH103531 and R01MH116901 to JA, T32NS099042 and 1F31MH125510 to EEB, T32GM007635 to BD and from the Brain & Behavior Research Foundation (NARSAD24847 to JA).

## References

Aoto, J., Martinelli, D. C., Malenka, R. C., Tabuchi, K., & Südhof, T. C. (2013). Presynaptic neurexin-3 alternative splicing trans-synaptically controls postsynaptic AMPA receptor trafficking. Cell, 154(1), 75–88. https://doi.org/10.1016/j.cell.2013.05.060

Baimel, C., Jang, E., Scudder, S. L., Manoocheri, K., & Carter, A. G. (2022). Hippocampal-evoked inhibition of cholinergic interneurons in the nucleus accumbens. Cell Reports, 40(1), 111042. https://doi.org/10.1016/j.celrep.2022.111042

Baimel, C., McGarry, L. M., & Carter, A. G. (2019). The Projection Targets of Medium Spiny Neurons Govern Cocaine-Evoked Synaptic Plasticity in the Nucleus Accumbens. Cell Reports, 28(9), 2256-2263.e3. https://doi.org/10.1016/j.celrep.2019.07.074

Becker, J. B. (2016). Sex differences in addiction. Dialogues in Clinical Neuroscience, 18(4), 395–402.

Beier, K. T., Kim, C. K., Hoerbelt, P., Hung, L. W., Heifets, B. D., DeLoach, K. E., Mosca, T. J., Neuner, S., Deisseroth, K., Luo, L., & Malenka, R. C. (2017). Rabies screen reveals GPe control of cocaine-triggered plasticity. Nature, 549(7672), 345–350. https://doi.org/10.1038/nature23888

Bekkers, J. M., & Clements, J. D. (1999). Quantal amplitude and quantal variance of strontium-induced asynchronous EPSCs in rat dentate granule neurons. The Journal of Physiology, 516(1), 227–248. https://doi.org/10.1111/j.1469-7793.1999.227aa.x

Bock, R., Shin, J. H., Kaplan, A. R., Dobi, A., Markey, E., Kramer, P. F., Gremel, C. M., Christensen, C. H., Adrover, M. F., & Alvarez, V. A. (2013). Strengthening the accumbal indirect pathway promotes resilience to compulsive cocaine use. Nature Neuroscience, 16(5), 632–638. https://doi.org/10.1038/nn.3369

Böhm, C., Peng, Y., Maier, N., Winterer, J., Poulet, J. F. A., Geiger, J. R. P., & Schmitz, D. (2015). Functional Diversity of Subicular Principal Cells during Hippocampal Ripples. The Journal of Neuroscience, 35(40), 13608–13618. https://doi.org/10.1523/JNEUROSCI.5034-14.2015

Bossert, J. M., Adhikary, S., St. Laurent, R., Marchant, N. J., Wang, H.-L., Morales, M., & Shaham, Y. (2016). Role of projections from ventral subiculum to nucleus accumbens shell in context-induced reinstatement of heroin seeking in rats. Psychopharmacology, 233(10), 1991–2004. https://doi.org/10.1007/s00213-015-4060-5

Boxer, E. E., Seng, C., Lukacsovich, D., Kim, J., Schwartz, S., Kennedy, M. J., Földy, C., & Aoto, J. (2021). Neurexin-3 defines synapse- and sex-dependent diversity of GABAergic inhibition in ventral subiculum. Cell Reports, 37(10), 110098. https://doi.org/10.1016/j.celrep.2021.110098

Britt, J. P., Benaliouad, F., McDevitt, R. A., Stuber, G. D., Wise, R. A., & Bonci, A. (2012). Synaptic and Behavioral Profile of Multiple Glutamatergic Inputs to the Nucleus Accumbens. Neuron, 76(4), 790–803. https://doi.org/10.1016/j.neuron.2012.09.040

Cahill, E., Pascoli, V., Trifilieff, P., Savoldi, D., Kappès, V., Lüscher, C., Caboche, J., & Vanhoutte, P. (2014). D1R/GluN1 complexes in the striatum integrate dopamine and glutamate signalling to control synaptic plasticity and cocaine-induced responses. Molecular Psychiatry, 19(12), Article 12. https://doi.org/10.1038/mp.2014.73

Castro, D. C., & Bruchas, M. R. (2019). A Motivational and Neuropeptidergic Hub: Anatomical and Functional Diversity within the Nucleus Accumbens Shell. Neuron, 102(3), 529–552. https://doi.org/10.1016/j.neuron.2019.03.003

Cembrowski, M. S., Phillips, M. G., DiLisio, S. F., Shields, B. C., Winnubst, J., Chandrashekar, J., Bas, E., & Spruston, N. (2018). Dissociable Structural and Functional Hippocampal Outputs via Distinct Subiculum Cell Classes. Cell, 173(5), 1280-1292.e18. https://doi.org/10.1016/j.cell.2018.03.031

Ciocchi, S., Passecker, J., Malagon-Vina, H., Mikus, N., & Klausberger, T. (2015). Selective information routing by ventral hippocampal CA1 projection neurons. Science, 348(6234), 560–563. https://doi.org/10.1126/science.aaa3245

Cole, S., Robinson, M. J. F., & Berridge, K. C. (2018). Optogenetic self-stimulation in the nucleus accumbens: D1 reward versus D2 ambivalence. PloS One, 13(11), e0207694. https://doi.org/10.1371/journal.pone.0207694

Dunn, A. R., Neuner, S. M., Ding, S., Hope, K. A., O’Connell, K. M. S., & Kaczorowski, C. C. (2018). Cell-Type-Specific Changes in Intrinsic Excitability in the Subiculum following Learning and Exposure to Novel Environmental Contexts. ENeuro, 5(6). https://doi.org/10.1523/ENEURO.0484-18.2018

Fuccillo, M. V., Földy, C., Gökce, Ö., Rothwell, P. E., Sun, G. L., Malenka, R. C., & Südhof, T. C. (2015). Single-Cell mRNA Profiling Reveals Cell-Type-Specific Expression of Neurexin Isoforms. Neuron, 87(2), 326–340. https://doi.org/10.1016/j.neuron.2015.06.028

Glangetas, C., Fois, G. R., Jalabert, M., Lecca, S., Valentinova, K., Meye, F. J., Diana, M., Faure, P., Mameli, M., Caille, S., & Georges, F. (2015). Ventral Subiculum Stimulation Promotes Persistent Hyperactivity of Dopamine Neurons and Facilitates Behavioral Effects of Cocaine. Cell Reports, 13(10), 2287–2296. https://doi.org/10.1016/j.celrep.2015.10.076

Gong, S., Fayette, N., Heinsbroek, J. A., & Ford, C. P. (2021). Cocaine shifts dopamine D2 receptor sensitivity to gate conditioned behaviors. Neuron. https://doi.org/10.1016/j.neuron.2021.08.012

Goto, Y., & Grace, A. A. (2005). Dopamine-Dependent Interactions between Limbic and Prefrontal Cortical Plasticity in the Nucleus Accumbens: Disruption by Cocaine Sensitization. Neuron, 47(2), 255–266. https://doi.org/10.1016/j.neuron.2005.06.017

Grace, A. A. (2010). Dopamine system dysregulation by the ventral subiculum as the common pathophysiological basis for schizophrenia psychosis, psychostimulant abuse, and stress. Neurotoxicity Research, 18(3–4), 367–376. https://doi.org/10.1007/s12640-010-9154-6

Gradinaru, V., Zhang, F., Ramakrishnan, C., Mattis, J., Prakash, R., Diester, I., Goshen, I., Thompson, K. R., & Deisseroth, K. (2010). Molecular and cellular approaches for diversifying and extending optogenetics. Cell, 141(1), 154–165. https://doi.org/10.1016/j.cell.2010.02.037

Graves, A. R., Moore, S. J., Bloss, E. B., Mensh, B. D., Kath, W. L., & Spruston, N. (2012). Hippocampal Pyramidal Neurons Comprise Two Distinct Cell Types that Are Countermodulated by Metabotropic Receptors. Neuron, 76(4), 776–789. https://doi.org/10.1016/j.neuron.2012.09.036

Groenewegen, H. J., Mulder, A. B., Beijer, A. V. J., Wright, C. I., Lopes da Silva, F. H., & Pennartz, C. M. A. (1999). Hippocampal and amygdaloid interactions in the nucleus accumbens. Psychobiology, 27(2), 149–164. https://doi.org/10.3758/BF03332111

Håkansson, K., Galdi, S., Hendrick, J., Snyder, G., Greengard, P., & Fisone, G. (2006). Regulation of phosphorylation of the GluR1 AMPA receptor by dopamine D2 receptors. Journal of Neurochemistry, 96(2), 482–488. https://doi.org/10.1111/j.1471-4159.2005.03558.x

Iino, Y., Sawada, T., Yamaguchi, K., Tajiri, M., Ishii, S., Kasai, H., & Yagishita, S. (2020). Dopamine D2 receptors in discrimination learning and spine enlargement. Nature, 579(7800), Article 7800. https://doi.org/10.1038/s41586-020-2115-1

Inbar, K., Levi, L. A., & Kupchik, Y. M. (2022). Cocaine induces input and cell-type-specific synaptic plasticity in ventral pallidum-projecting nucleus accumbens medium spiny neurons. Neuropsychopharmacology, 1–12. https://doi.org/10.1038/s41386-022-01285-6

Jia, Z., Agopyan, N., Miu, P., Xiong, Z., Henderson, J., Gerlai, R., Taverna, F. A., Velumian, A., MacDonald, J., Carlen, P., Abramow-Newerly, W., & Roder, J. (1996). Enhanced LTP in mice deficient in the AMPA receptor GluR2. Neuron, 17(5), 945–956. https://doi.org/10.1016/s0896-6273(00)80225-1

Kim, Y., & Spruston, N. (2012). Target-specific output patterns are predicted by the distribution of regular-spiking and bursting pyramidal neurons in the subiculum. Hippocampus, 22(4), 693–706. https://doi.org/10.1002/hipo.20931

Lamsa, K. P., Heeroma, J. H., Somogyi, P., Rusakov, D. A., & Kullmann, D. M. (2007). Anti-Hebbian Long-Term Potentiation in the Hippocampal Feedback Inhibitory Circuit. Science (New York, N.Y.), 315(5816), 1262–1266. https://doi.org/10.1126/science.1137450

Lee, S., Lee, C., Woo, C., Kang, S. J., & Shin, K. S. (2019). Chronic social defeat stress increases burst firing of nucleus accumbens-projecting ventral subicular neurons in stress-susceptible mice. Biochemical and Biophysical Research Communications, 515(3), 468–473. https://doi.org/10.1016/j.bbrc.2019.05.180

LeGates, T. A., Kvarta, M. D., Tooley, J. R., Francis, T. C., Lobo, M. K., Creed, M. C., & Thompson, S. M. (2018). Reward behaviour is regulated by the strength of hippocampus–nucleus accumbens synapses. Nature, 564(7735), Article 7735. https://doi.org/10.1038/s41586-018-0740-8

Li, Z., Chen, Z., Fan, G., Li, A., Yuan, J., & Xu, T. (2018). Cell-Type-Specific Afferent Innervation of the Nucleus Accumbens Core and Shell. Frontiers in Neuroanatomy, 12. https://www.frontiersin.org/articles/10.3389/fnana.2018.00084

Lindenbach, D., Vacca, G., Ahn, S., Seamans, J. K., & Phillips, A. G. (2022). Optogenetic modulation of glutamatergic afferents from the ventral subiculum to the nucleus accumbens: Effects on dopamine function, response vigor and locomotor activity. Behavioural Brain Research, 434, 114028. https://doi.org/10.1016/j.bbr.2022.114028

Liu, X.-Y., Chu, X.-P., Mao, L.-M., Wang, M., Lan, H.-X., Li, M.-H., Zhang, G.-C., Parelkar, N. K., Fibuch, E. E., Haines, M., Neve, K. A., Liu, F., Xiong, Z.-G., & Wang, J. Q. (2006). Modulation of D2R-NR2B interactions in response to cocaine. Neuron, 52(5), 897–909. https://doi.org/10.1016/j.neuron.2006.10.011

Liu, Z., Le, Q., Lv, Y., Chen, X., Cui, J., Zhou, Y., Cheng, D., Ma, C., Su, X., Xiao, L., Yang, R., Zhang, J., Ma, L., & Liu, X. (2022). A distinct D1-MSN subpopulation down-regulates dopamine to promote negative emotional state. Cell Research, 32(2), Article 2. https://doi.org/10.1038/s41422-021-00588-5

Lobo, M. K., Covington, H. E., Chaudhury, D., Friedman, A. K., Sun, H., Damez-Werno, D., Dietz, D. M., Zaman, S., Koo, J. W., Kennedy, P. J., Mouzon, E., Mogri, M., Neve, R. L., Deisseroth, K., Han, M.-H., & Nestler, E. J. (2010). Cell Type–Specific Loss of BDNF Signaling Mimics Optogenetic Control of Cocaine Reward. Science, 330(6002), 385–390. https://doi.org/10.1126/science.1188472

Lobo, M. K., & Nestler, E. (2011). The Striatal Balancing Act in Drug Addiction: Distinct Roles of Direct and Indirect Pathway Medium Spiny Neurons. Frontiers in Neuroanatomy, 5, 41. https://doi.org/10.3389/fnana.2011.00041

Lopes da Silva, F. H., Arnolds, D. E., & Neijt, H. C. (1984). A functional link between the limbic cortex and ventral striatum: Physiology of the subiculum accumbens pathway. Experimental Brain Research, 55(2), 205–214. https://doi.org/10.1007/BF00237271

MacAskill, A. F., Cassel, J. M., & Carter, A. G. (2014). Cocaine exposure reorganizes cell type- and input-specific connectivity in the nucleus accumbens. Nature Neuroscience, 17(9), 1198–1207. https://doi.org/10.1038/nn.3783

Mameli, M., Bellone, C., Brown, M. T. C., & Lüscher, C. (2011). Cocaine inverts rules for synaptic plasticity of glutamate transmission in the ventral tegmental area. Nature Neuroscience, 14(4), 414–416. https://doi.org/10.1038/nn.2763

Marchant, N. J., Campbell, E. J., Whitaker, L. R., Harvey, B. K., Kaganovsky, K., Adhikary, S., Hope, B. T., Heins, R. C., Prisinzano, T. E., Vardy, E., Bonci, A., Bossert, J. M., & Shaham, Y. (2016). Role of Ventral Subiculum in Context-Induced Relapse to Alcohol Seeking after Punishment-Imposed Abstinence. Journal of Neuroscience, 36(11), 3281–3294. https://doi.org/10.1523/JNEUROSCI.4299-15.2016

Meredith, G. E., Baldo, B. A., Andrezjewski, M. E., & Kelley, A. E. (2008). The structural basis for mapping behavior onto the ventral striatum and its subdivisions. Brain Structure and Function, 213(1), 17–27. https://doi.org/10.1007/s00429-008-0175-3

Naber, P. a., & Witter, M. p. (1998). Subicular efferents are organized mostly as parallel projections: A double-labeling, retrograde-tracing study in the rat. Journal of Comparative Neurology, 393(3), 284–297. https://doi.org/10.1002/(SICI)1096-9861(19980413)393:3<284::AID-CNE2>3.0.CO;2-Y

Okuyama, T., Kitamura, T., Roy, D. S., Itohara, S., & Tonegawa, S. (2016). Ventral CA1 neurons store social memory. Science, 353(6307), 1536–1541. https://doi.org/10.1126/science.aaf7003

Park, P., Kang, H., Georgiou, J., Zhuo, M., Kaang, B.-K., & Collingridge, G. L. (2021). Further evidence that CP-AMPARs are critically involved in synaptic tag and capture at hippocampal CA1 synapses. Molecular Brain, 14(1), 26. https://doi.org/10.1186/s13041-021-00737-2

Pascoli, V., Terrier, J., Espallergues, J., Valjent, E., O’Connor, E. C., & Lüscher, C. (2014). Contrasting forms of cocaine-evoked plasticity control components of relapse. Nature, 509(7501), 459–464. https://doi.org/10.1038/nature13257

Pawlak, V., & Kerr, J. N. D. (2008). Dopamine Receptor Activation Is Required for Corticostriatal Spike-Timing-Dependent Plasticity. Journal of Neuroscience, 28(10), 2435–2446. https://doi.org/10.1523/JNEUROSCI.4402-07.2008

Preston, C. J., Brown, K. A., & Wagner, J. J. (2019). Cocaine conditioning induces persisting changes in ventral hippocampus synaptic transmission, long-term potentiation, and radial arm maze performance in the mouse. Neuropharmacology, 150, 27–37. https://doi.org/10.1016/j.neuropharm.2019.02.033

Roggenhofer, E., Fidzinski, P., Shor, O., & Behr, J. (2013). Reduced Threshold for Induction of LTP by Activation of Dopamine D1/D5 Receptors at Hippocampal CA1–Subiculum Synapses. PLOS ONE, 8(4), e62520. https://doi.org/10.1371/journal.pone.0062520

Rothwell, P. E., Fuccillo, M. V., Maxeiner, S., Hayton, S. J., Gokce, O., Lim, B. K., Fowler, S. C., Malenka, R. C., & Südhof, T. C. (2014). Autism-Associated Neuroligin-3 Mutations Commonly Impair Striatal Circuits to Boost Repetitive Behaviors. Cell, 158(1), 198–212. https://doi.org/10.1016/j.cell.2014.04.045

Schotanus, S. M., & Chergui, K. (2008). Dopamine D1 receptors and group I metabotropic glutamate receptors contribute to the induction of long-term potentiation in the nucleus accumbens. Neuropharmacology, 54(5), 837–844. https://doi.org/10.1016/j.neuropharm.2007.12.012

Scudder, S. L., Baimel, C., Macdonald, E. E., & Carter, A. G. (2018). Hippocampal-Evoked Feedforward Inhibition in the Nucleus Accumbens. The Journal of Neuroscience: The Official Journal of the Society for Neuroscience, 38(42), 9091–9104. https://doi.org/10.1523/JNEUROSCI.1971-18.2018

Sinnen, B. L., Bowen, A. B., Forte, J. S., Hiester, B. G., Crosby, K. C., Gibson, E. S., Dell’Acqua, M. L., & Kennedy, M. J. (2017). Optogenetic Control of Synaptic Composition and Function. Neuron, 93(3), 646-660.e5. https://doi.org/10.1016/j.neuron.2016.12.037

Sjulson, L., Peyrache, A., Cumpelik, A., Cassataro, D., & Buzsáki, G. (2018). Cocaine Place Conditioning Strengthens Location-Specific Hippocampal Coupling to the Nucleus Accumbens. Neuron, 98(5), 926-934.e5. https://doi.org/10.1016/j.neuron.2018.04.015

Soares-Cunha, C., Coimbra, B., Sousa, N., & Rodrigues, A. J. (2016). Reappraising striatal D1- and D2-neurons in reward and aversion. Neuroscience & Biobehavioral Reviews, 68, 370–386. https://doi.org/10.1016/j.neubiorev.2016.05.021

Soares-Cunha, C., Domingues, A. V., Correia, R., Coimbra, B., Vieitas-Gaspar, N., de Vasconcelos, N. A. P., Pinto, L., Sousa, N., & Rodrigues, A. J. (2022). Distinct role of nucleus accumbens D2-MSN projections to ventral pallidum in different phases of motivated behavior. Cell Reports, 38(7), 110380. https://doi.org/10.1016/j.celrep.2022.110380

Staff, N. P., Jung, H.-Y., Thiagarajan, T., Yao, M., & Spruston, N. (2000). Resting and Active Properties of Pyramidal Neurons in Subiculum and CA1 of Rat Hippocampus. Journal of Neurophysiology, 84(5), 2398–2408. https://doi.org/10.1152/jn.2000.84.5.2398

Südhof, T. C. (2017). Synaptic Neurexin Complexes: A Molecular Code for the Logic of Neural Circuits. Cell, 171(4), 745–769. https://doi.org/10.1016/j.cell.2017.10.024

Terrier, J., Lüscher, C., & Pascoli, V. (2016). Cell-Type Specific Insertion of GluA2-Lacking AMPARs with Cocaine Exposure Leading to Sensitization, Cue-Induced Seeking, and Incubation of Craving. Neuropsychopharmacology, 41(7), 1779–1789. https://doi.org/10.1038/npp.2015.345

Williams, E. S., Manning, C. E., Eagle, A. L., Swift-Gallant, A., Duque-Wilckens, N., Chinnusamy, S., Moeser, A., Jordan, C., Leinninger, G., & Robison, A. J. (2020). Androgen-Dependent Excitability of Mouse Ventral Hippocampal Afferents to Nucleus Accumbens Underlies Sex-Specific Susceptibility to Stress. Biological Psychiatry, 87(6), 492–501. https://doi.org/10.1016/j.biopsych.2019.08.006

Wozny, C., Maier, N., Schmitz, D., & Behr, J. (2008). Two different forms of long-term potentiation at CA1–subiculum synapses. The Journal of Physiology, 586(Pt 11), 2725–2734. https://doi.org/10.1113/jphysiol.2007.149203

Yang, H., de Jong, J. W., Tak, Y., Peck, J., Bateup, H. S., & Lammel, S. (2018). Nucleus Accumbens Subnuclei Regulate Motivated Behavior via Direct Inhibition and Disinhibition of VTA Dopamine Subpopulations. Neuron, 97(2), 434-449.e4. https://doi.org/10.1016/j.neuron.2017.12.022

Yao, Y., Gao, G., Liu, K., Shi, X., Cheng, M., Xiong, Y., & Song, S. (2021). Projections from D2 Neurons in Different Subregions of Nucleus Accumbens Shell to Ventral Pallidum Play Distinct Roles in Reward and Aversion. Neuroscience Bulletin, 37(5), 623–640. https://doi.org/10.1007/s12264-021-00632-9

Yu, J., Yan, Y., Li, K.-L., Wang, Y., Huang, Y. H., Urban, N. N., Nestler, E. J., Schlüter, O. M., & Dong, Y. (2017). Nucleus accumbens feedforward inhibition circuit promotes cocaine self-administration. Proceedings of the National Academy of Sciences, 114(41), E8750–E8759. https://doi.org/10.1073/pnas.1707822114

Zahm, D. S. (1992). An electron microscopic morphometric comparison of tyrosine hydroxylase immunoreactive innervation in the neostriatum and the nucleus accumbens core and shell. Brain Research, 575(2), 341–346. https://doi.org/10.1016/0006-8993(92)90102-F

Zhou, Y., Zhu, H., Liu, Z., Chen, X., Su, X., Ma, C., Tian, Z., Huang, B., Yan, E., Liu, X., & Ma, L. (2019). A ventral CA1 to nucleus accumbens core engram circuit mediates conditioned place preference for cocaine. Nature Neuroscience, 22(12), 1986–1999. https://doi.org/10.1038/s41593-019-0524-y

